# A mycobacterial DNA extraction protocol designed for resource limited settings generates high quality whole genome sequencing

**DOI:** 10.1101/2024.05.31.596815

**Authors:** Caitlin Percy, Ilinca Memelis, Thomas Edwards, Adam P. Roberts, Giancarlo Biagini, Daire Cantillon

## Abstract

Mycobacteria are major global human pathogens and include *Mycobacterium tuberculosis*, the causative agent of tuberculosis, and *M. abscessus*, an emerging multidrug resistant pathogen.

*M. abscessus* affects people with structural lung disease and those who are immunocompromised, most commonly causing pulmonary disease but also disseminated infections in the central nervous system and skin. High quality whole genome sequencing is essential to research mycobacterial epidemiology, pathogenesis and antimicrobial resistance. However, current DNA extraction protocols are time consuming, use toxic chemicals, require cold chain storage for certain reagents and can often result in poor quality, degraded DNA that directly impacts whole genome sequencing outputs. This is a particular challenge in low-income settings.

Here, we report a novel optimised DNA extraction workflow for *M. tuberculosis* and *M. abscessus* that invariably generates high quality Illumina short read sequencing data. We evaluated input culture CFU and physical cell disruption times. DNA quantity was determined using a Qubit fluorometer system with DNA integrity assessed using the Agilent TapeStation platform.

We showed that this protocol facilitated complete genome assemblies of *M. abscessus* and *M. tuberculosis* reference strains. There is no requirement for cold chain transport or storage of reagents, solvent extractions, or boiling to heat inactivate cultures, and the method does not require surfactant chemicals such as cetyltrimethylammonium bromide (CTAB).

## Introduction

Tuberculosis (TB), caused by the bacterial pathogen *Mycobacterium tuberculosis*, is a respiratory infectious disease with 7.5 million new cases recorded in 2022, the highest ever reported since the World Health Organisation (WHO) began global TB monitoring in 1995.

Despite TB being largely preventable and curable, it is now the leading cause of death due to a single infectious agent globally with 1.5 million deaths annually. Drug resistance is a formidable challenge with an estimated 410,000 people developing rifampicin (RIF) resistant or multidrug resistance TB (RR/MDR-TB) but only 40% of these accessing diagnosis and treatment (World Health Organisation 2023). RIF resistant TB has also become a marker for multi-drug resistant

(MDR-TB) due to there being limited rifampicin mono-resistant strains of TB; 85-90% of RIF resistant TB is also resistant to isoniazid (INH) (Pang, Lu et al. 2013, Malenfant and Brewer 2021). Globally, the number of people diagnosed with pulmonary TB and subsequently tested for RIF resistance was 71% in 2020, increasing from 61% in 2019. However, as a repercussion of the COVID19 pandemic, the case detection rate of new TB infections fell by 18% between 2019 and 2020 (World Health Organisation 2022). The global increase in drug resistance in TB highlights the need for wider application of next generation sequencing, and more efficient, user friendly, reproducible DNA extraction protocols to support these platforms.

Non-tuberculous mycobacteria (NTM) are an emerging global health threat. Broadly, NTMs are classified into fast growers, including *Mycobacterium abscessus* (*M. abscessus*), and slow growers such as *M. avium* (Johansen, Herrmann et al. 2020). *M. abscessus* infections result in prolonged treatment times (12-18 months) of multi-drug therapy with a treatment success rate of just 33-45% (Diel, Ringshausen et al. 2017, Haworth, Banks et al. 2017). Drugs used to treat *M. tuberculosis* do not work for *M. abscessus* as it is intrinsically resistant to all first line TB antibiotics (Cantillon, Goff et al. 2022). Whole genome sequencing to detect markers of antimicrobial resistance is not currently used to guide treatment as is the case for *M. tuberculosis*, although there is impetus for this to be developed (Dohal, Porvaznik et al. 2021). In addition, retrospective studies in certain LMICs show that NTM infection has been mis-identified as MDR TB, with up to 30% of cases in Iran and 15% in Zimbabwe reporting this misdiagnosis (Chanda-Kapata, Kapata et al. 2015, Shahraki, Heidarieh et al. 2015).

Currently there is no universally adopted methodology for DNA extraction for mycobacteria in resource limited settings, however the cetyltrimethylammonium bromide (CTAB) method is widely implemented. The original CTAB method was developed as a plant DNA extraction method but was subsequently applied to mycobacteria (Doyle and Doyle 1990). Briefly, the CTAB method for mycobacteria involves lysis with lysozyme and proteinase K, CTAB and other chemicals to precipitate the DNA followed by several wash steps (Jagatia and Cantillon 2021). CTAB can be advantageous due to the low cost and PCR inhibitor removal. However, the method makes use of toxic chemical such as CTAB and SDS, is time consuming and often results in short, fragmented, poor-quality DNA that fails sequencing (McNerney, Clark et al. 2017). Therefore, due to the lack of universally adopted protocol for DNA extraction in mycobacteria and the methods that do exist having high sequencing failure rates, there is a need to develop a universally adoptable DNA extraction method in mycobacteria.

This study set out to develop a user friendly, rapid DNA extraction method suitable for low resource settings. Typically, culture samples are heat inactivated by boiling or heat blocks. Due to safety risks with this approach, as well as non-uniform heating and killing of samples (Bemer-Melchior and Drugeon 1999), we validated 70% ethanol inactivation of *M. tuberculosis* instead. We demonstrate the method facilitates high quality short read whole genome sequencing of two medically important mycobacteria, *M. tuberculosis* and *M. abscessus*.

## Materials and Methods

### Strains and culture conditions

*M. abscessus* reference strain ATCC19977 and *M. tuberculosis* reference strain H37Rv were cultured in Middlebrook 7H9 broth (Sigma Aldrich, St Louis, MO, USA) supplemented with oleic acid albumin dextrose catalase (OADC; 10% v/v) and Tween 80 (0·05% v/v; Sigma Aldrich, St Louis, MO, USA) at 37°C. For solid media culture, 7H10 agar (Sigma Aldrich, St Louis, MO, USA) was supplemented with OADC (10% v/v) and glycerol (0.5% v/v). Optical density (O.D.) was determined by using a spectrophotometer at 600 nm. *M. tuberculosis* laboratory work was carried out at BSL3, while *M. abscessus* was worked with at BSL2.

### Optimisation of inactivation of M. tuberculosis

Log phase *M. tuberculosis* H37Rv cultures were O.D. adjusted to 1.0 in 1 mL aliquots in 1.5 mL Eppendorf tubes, corresponding to ∼1 x 10^8^ CFU/mL per tube. These were centrifuged at 12,000 rcf for 1 minute, supernatant discarded and pellets resuspended in 1.5 mL 70% ethanol made fresh on the day. Pellets were incubated for 10, 20, 40 and 60 minutes with mixing by inversion five times half ways through each timepoint. To harvest each timepoint, tubes were centrifuged at 12,000 rcf for 1 minute, 70% ethanol supernatant discarded, and pellets resuspended in 1 mL PBS and immediately plated out on to 7H10 agar, allowed dry and incubated for six weeks at 37°C. The OD 1.0 adjusted inoculum was serially diluted in PBS and also plated out and incubated for four weeks at 37°C. CFU/mL was determined using the following formula:

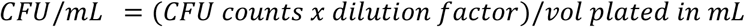

At least a seven-log (99.99999%) reduction was required to consider *M. tuberculosis* inactivated.

### DNA extraction from M. tuberculosis and M. abscessus

DNeasy PowerBiofilm DNA extraction kit (Qiagen, UK) was used to extract genomic DNA. *M. abscessus* and ethanol inactivated *M. tuberculosis* 1 mL O.D. 1.0 (∼10^8^ CFU/mL) culture aliquots were centrifuged at 12,000 rcf for 1 minute, supernatant discarded, resuspended in Solution MBL, and transferred to the bead tubes supplied with the kit. Various bead beating times of 15 seconds, 30 seconds, 45 seconds and 60 seconds at 10,000 rpm for both species and an additional 120 seconds for *M. tuberculosis* were evaluated in a Precellys ribolyser (Bertin Technologies, France). DNA was extracted by following the manufacturers’ instructions. A 20 minute incubation time of 70% ethanol for *M. tuberculosis* and bead beating time of 30 seconds was used for DNA extractions.

### Quality control

DNA extractions were quantified using Qubit 4 fluorometer with a double stranded DNA broad range quantification assay (ThermoFisher, UK) as per manufacturer’s instructions. The Tapestation 4150 (Agilent) microfluidic electrophoresis platform was used to assess DNA integrity through DNA integrity numbers (DINs).

### Whole Genome Sequencing

For llumina short read sequencing by MicrobesNG, libraries were prepared using the Nextera XT Library Prep Kit (Illumina, San Diego, USA) following manufacturer’s instructions. Libraries were sequenced on an Illumina NovaSeq 6000 (Illumina, San Diego, USA) using a 250 bp paired end read protocol. Reads were subsequently trimmed using Trimmomatic (v 0.30) (Bolger, Lohse et al. 2014) with a Q15 sliding window quality cutoff. SPAdes (v 3.7) (Bankevich, Nurk et al. 2012) was used for de novo assembly with contigs annotated using Prokka (v 1.11) (Seemann 2014).

### Genome assemblies with Bandage

Illumina short read data from MicrobesNG was used for genome assemblies with Bandage (v 0.8.1) via Pathosystems Resource Integration Center (PATRIC), a genomics-centric relational database and bioinformatics resource (Gillespie, Wattam et al. 2011) . Assembly used Unicycler (v 0.4.8) with one round of polishing by Pilon (Walker, Abeel et al. 2014). Quast (v 5.2.0) was used to generate Bandage assembly sequencing and quality control data tables, including longest and mean contigs (Mikheenko, Prjibelski et al. 2018) .

### Statistics

A paired parametric t test was used to assign statistical significance between groups in GraphPad Prism 10.1.1. Statistical significance was assigned based on the following p value ranges: No significance (ns) p > 0.05; ^*^ < 0.05; ^**^ < 0.01; ^***^ < 0.001; ^****^ < 0.0001.

## Results

### 70% ethanol effectively inactivates M. tuberculosis

The least amount of time for 70% ethanol treatment to reduce *M. tuberculosis* CFU/mL by at least seven-log was 10 minutes, the shortest timepoint tested, Table 1. This experiment was performed twice, with the data from both experiments in Table 1.

**Table 1:**
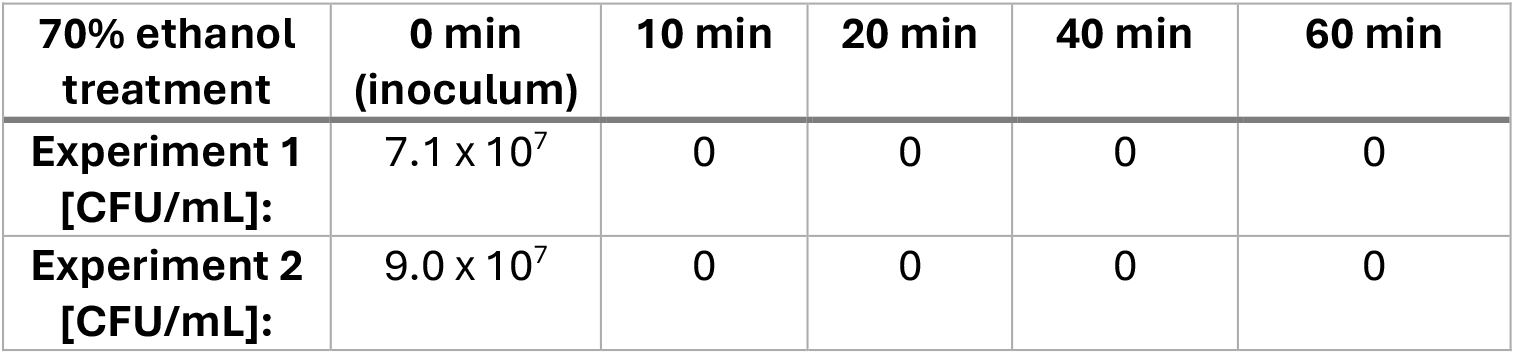
70% ethanol completely inhibits *M. tuberculosis* after 10 minutes by at least seven-log.

| 70% ethanol treatment | 0 min (inoculum) | 10 min | 20 min | 40 min | 60 min |
| --- | --- | --- | --- | --- | --- |
| Experiment 1 [CFU/mL]: | $7.1 \times 10^7$ | 0 | 0 | 0 | 0 |
| Experiment 2 [CFU/mL]: | $9.0 \times 10^7$ | 0 | 0 | 0 | 0 |

### Bead beating times dictate DNA yield and integrity

Cell lysis of mycobacteria is more challenging than other bacteria due to the robust, lipid-rich cell wall meaning enzymatic lysis can be inefficient. In this study, we employed mechanical lysis using bead beating. The PowerBiofilm kit states to bead beat at 3,200 rpm for 30 seconds; we hypothesised based on mycobacterial cell wall biology that this may be too low to break cells open as other studies bead beat at 10,000 rpm for mycobacteria successfully (Tailleux, Waddell et al. 2008, Bettencourt, Muller et al. 2020, Cantillon, Wroblewska et al. 2021). Bead beating of 10,000 rpm was then selected to go forward with and used at 15 seconds, 30 seconds, 45 seconds and 60 seconds for *M. abscessus* and *M. tuberculosis*, with an additional 120 second bead beating timepoint done for *M. tuberculosis*. For *M. abscessus*, bead beating of 45 and 60 seconds yielded significantly more DNA compared to 15 and 30 seconds, but there was a concomitant decrease in DIN, Figure 1.

**Figure 1:**
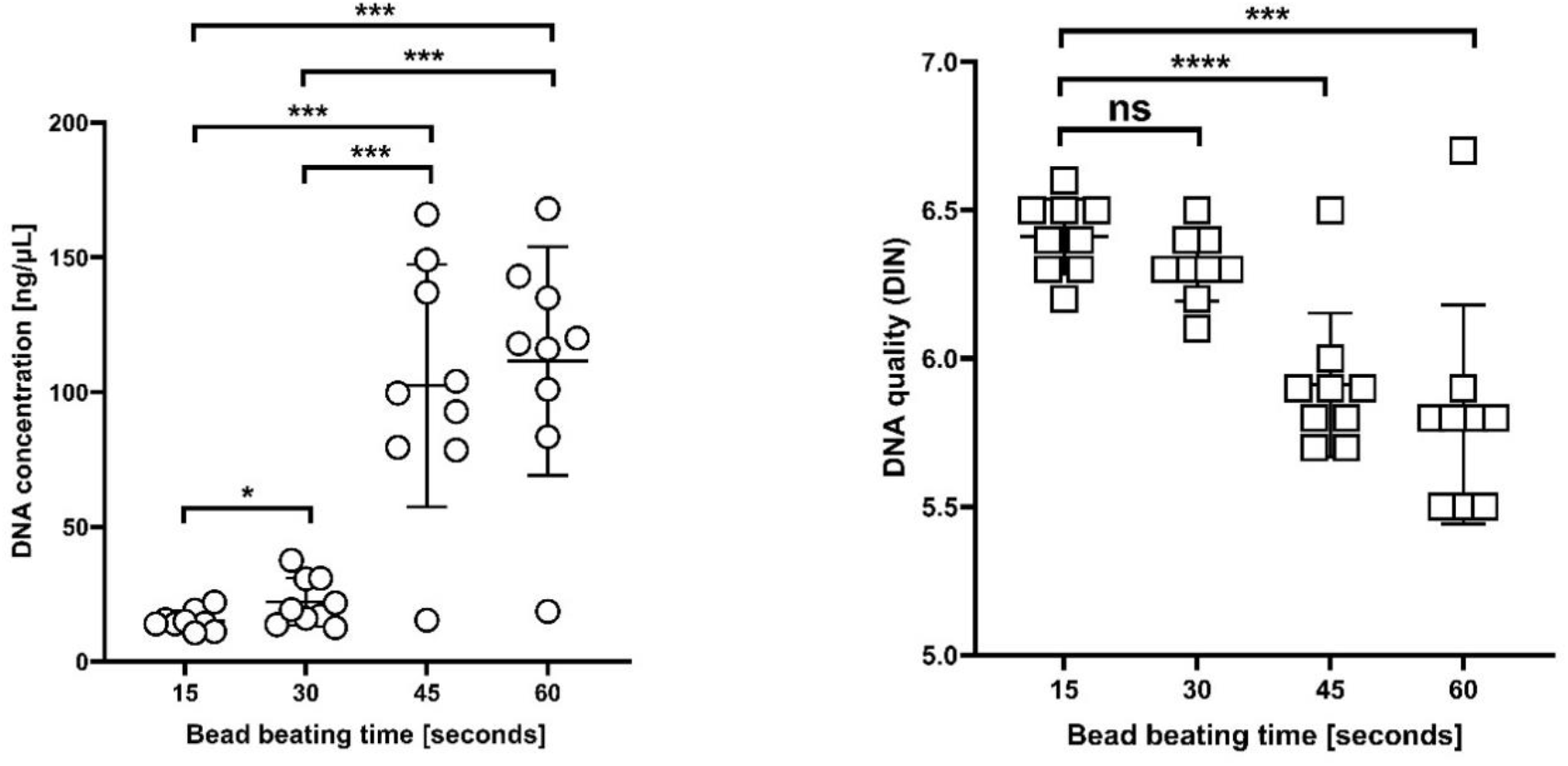
Bead beating times impact DNA yield and integrity in *M. abscessus*. DNA concentration was determined with Qubit, DIN determined with TapeStation. Increasing bead beating times increases DNA yield but reduces DIN. Data points are nine biological repeats pooled with standard deviation as error bars. No significance (ns) p > 0.05; ^*^ < 0.05; ^**^ < 0.01; ^***^ < 0.001; ^****^ < 0.0001.

For *M. tuberculosis*, the relationship between bead beating times and DIN was less clear compared to *M. abscessus*. There was no statistical significance between 15 seconds and 30, 45 or 60 seconds. Only 120 seconds timepoints showed a significantly decreased DIN compared to 15, 30, 45 and 60 second timepoints, suggesting that 120 seconds generates a DNA yield but that bead beating time significantly impacts on its integrity, Figure 2.

**Figure 2:**
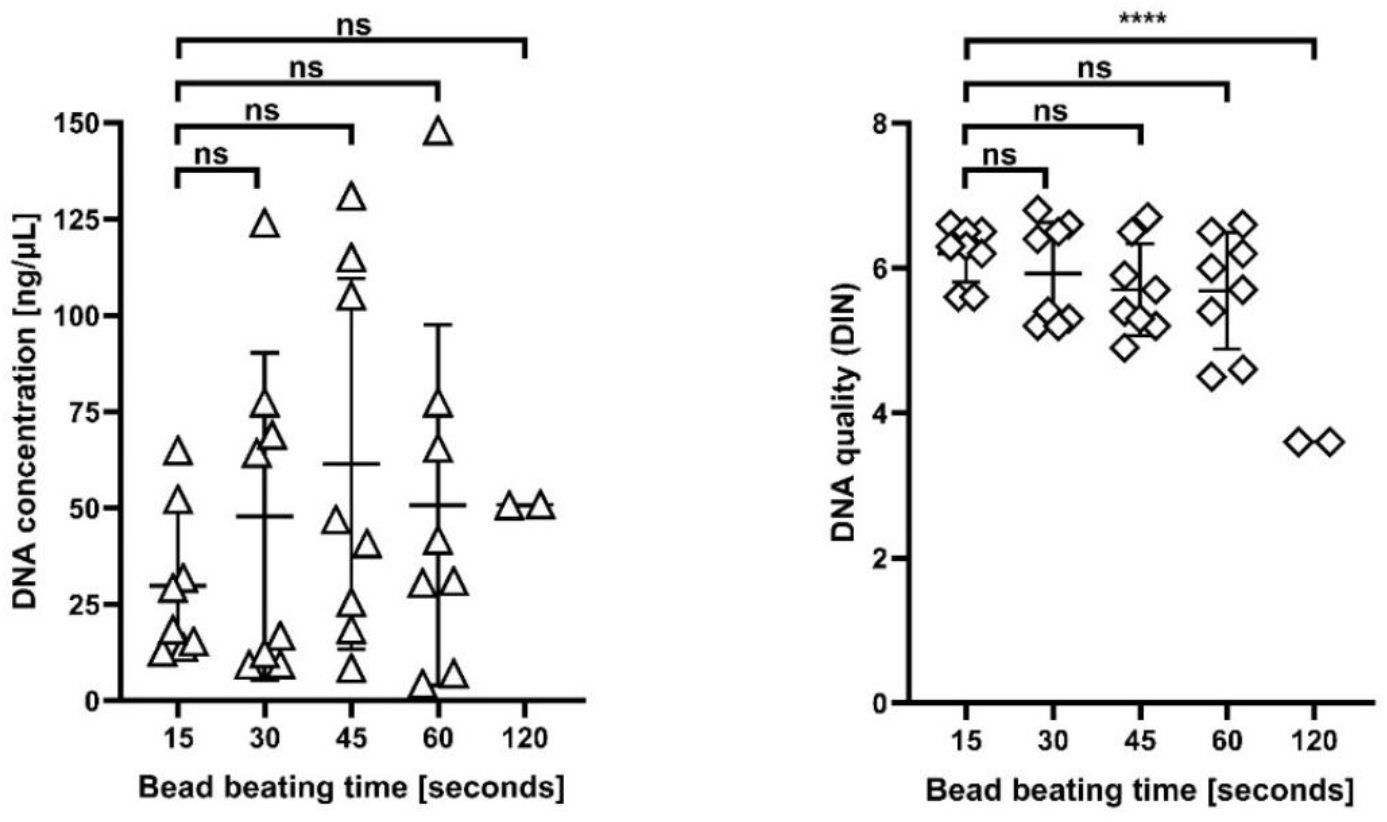
Bead beating times impact DNA yield and integrity in *M. tuberculosis*. DNA concentration was determined with Qubit, DIN determined with TapeStation. Increasing bead beating times increases DNA yield but reduces DIN. Data points are eight (except 120 seconds which is two) biological replicates pooled with standard deviation as error bars. No significance (ns) p > 0.05; ^*^ < 0.05; ^**^ < 0.01; ^***^ < 0.001; ^****^ < 0.0001.

### Minimum CFU providing measurable DNA is ∼10^7^ CFU/mL

DNA extraction methods in mycobacteria often involve a high input (>10^9^ CFU) of mycobacterial cells to extract enough DNA for sequencing. We wanted to establish how low the CFU input could be titrated down while still getting measurable DNA. To determine this, serial dilutions of *M. tuberculosis* and *M. abscessus* cultures were performed 1:10 in PBS, and each dilution extracted for DNA. The lower the amount of biomass inputted, the less DNA extracted, with some DNA still recoverable at O.D. 0.1 (∼10^7^ CFU/mL) for some samples (*M. abscessus* 6/8; *M. tuberculosis* 2/6) but this was <5 ng/µL while other O.D. 0.1 samples did not yield any detectable DNA (*M. abscessus* 2/8; *M. tuberculosis* n = 4/6). DIN values showed higher variation at lower O.D. samples, although were still relatively comparable to higher O.D. samples, Figure 2, Figure 3.

**Figure 3:**
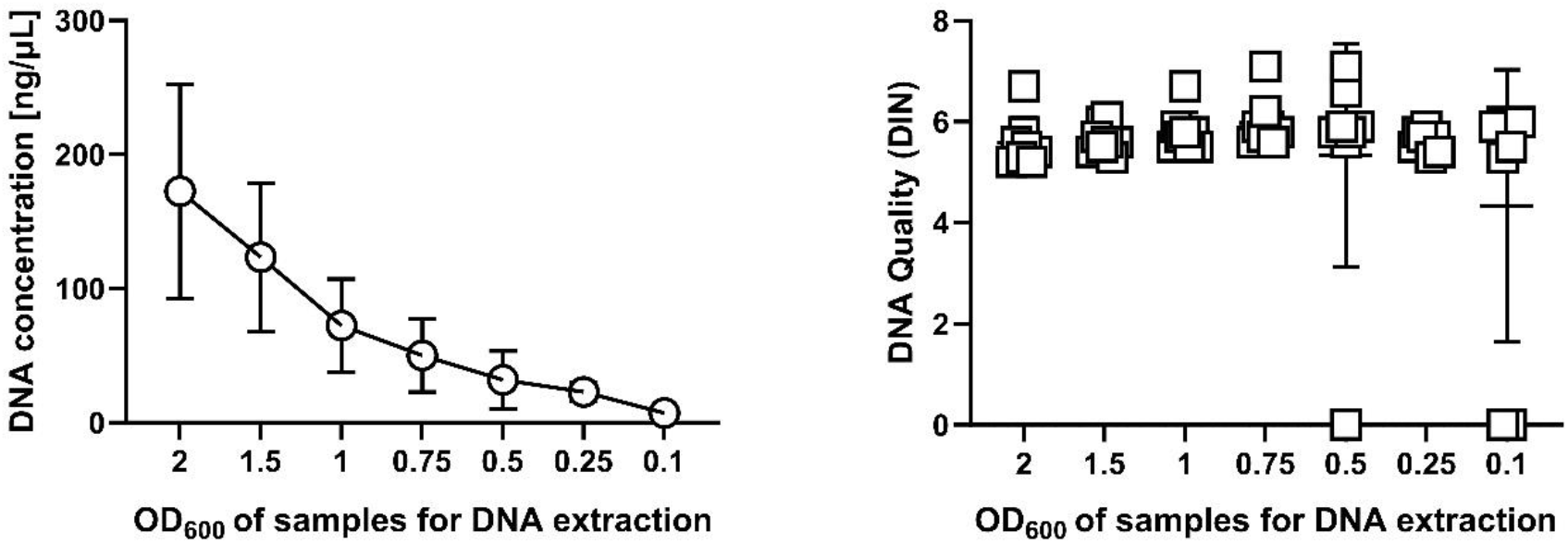
The limit of DNA extraction is O.D. 0.1 for *M. abscessus*. DNA concentration was determined with Qubit, DIN determined with TapeStation. Data points are eight biological repeats pooled with standard deviation as error bars.

**Figure 4:**
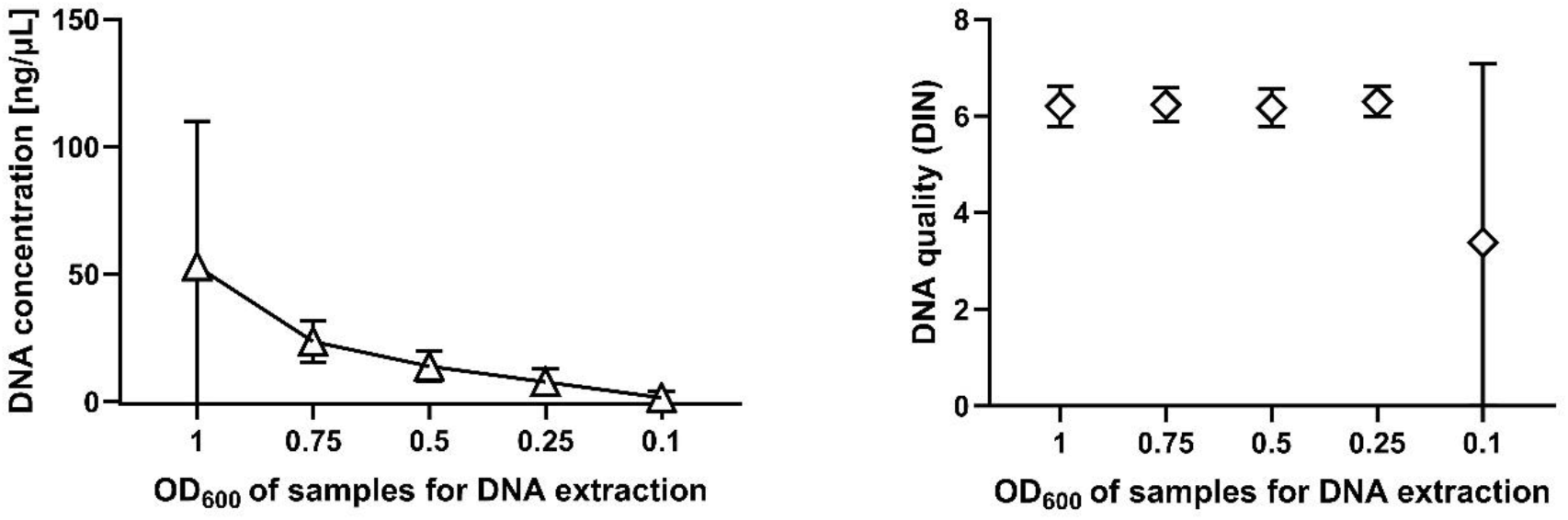
The limit of DNA extraction is O.D. 0.1 for *M. tuberculosis*. DNA concentration was determined with Qubit, DIN determined with TapeStation. Data points are six biological repeats pooled with standard deviation as error bars.

### Novel DNA extraction workflow facilitates high quality whole genome sequencing

DNA was extracted from 10^8^ CFU *M. tuberculosis* and *M. abscessus* and sent for whole genome sequencing by MicrobesNG using their Illumina NextSeq platform. Genomic DNA was extracted using 30 seconds bead beating and Qiagen DNeasy PowerBiofilm DNA extraction kit as per manufacturer’s instructions. High quality genome sequencing was obtained, Table 2.

**Table 2:** Key metrics of Illumina short read data.

| Species | Mean coverage | # contigs | # of reads | N50 |
| --- | --- | --- | --- | --- |
| *M. abscessus* | 86 | 14 | 928510 | 2798844 |
| *M. tuberculosis* | 184 | 61 | 1669534 | 147934 |

To visualise assemblies, short read data was processed using Bandage (via the PATRIC platform). For *M. abscessus*, Bandage shows long, continuous contigs with low fragmentation and simple graph topology with its longest contig 2,758,626 and a mean contig of 565,483.

Well-connected nodes with no breaks or branches suggest a complete and accurate sequence assembly. Bandage has also assembled a plasmid; ATCC19977 has been shown to contain a 23Kb mercury resistance plasmid (Ripoll, Pasek et al. 2009). For *M. tuberculosis*, the topology is more complex suggesting more fragmented assembly than *M. abscessus*. The more complete genome assembly of *M. abscessus* is reflected in that this strain had fewer contigs than *M. tuberculosis* (Figure 5).

**Figure 5:**
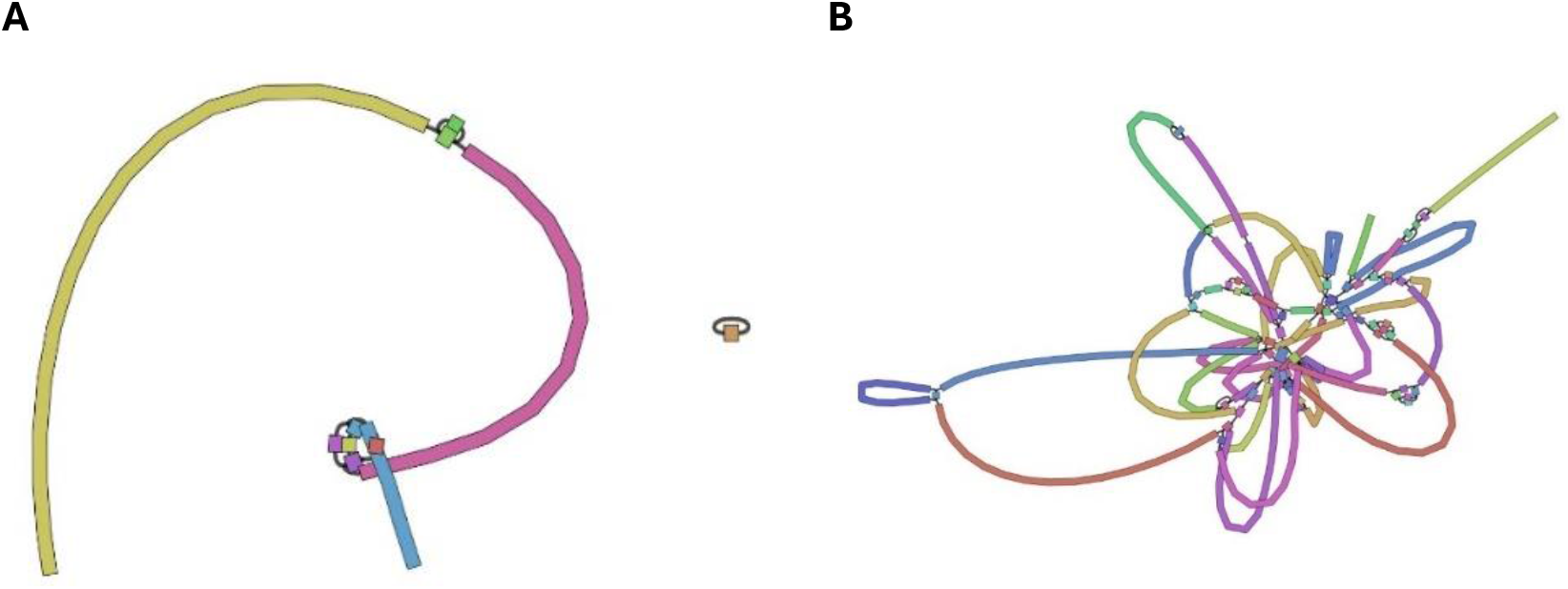
Bandage assemblies of A) *M. abscessus*, B) *M. tuberculosis* genomes. Short read Illumina data was analysed using Bandage

## Discussion

In order to extract DNA from *M. tuberculosis*, the culture must first be inactivated by at least seven-log in CFU for biosafety reasons to remove *M. tuberculosis* from BSL3 labs for DNA extractions (European Centre for Disease Prevention and Control 2018) either in house or for shipping to external providers of whole genome sequencing services. Heat killing of *M. tuberculosis* culture is the most widely adopted protocol of inactivation. Heating of a culture either involves a water bath, heat block or dry water bath. However, heat treatment has varying inactivation efficiencies, with some groups reporting uniform killing (Krugman 1975, Castro, Gonzalez et al. 2009), and others raising issues with kill consistencies (Billard-Pomares, Bleibtreu et al. 2019). Differences in temperature, duration and container type of heat inactivation has led to no uniform consensus being reached (Wang, Putri et al. 2021). Ethanol can be used as a chemical inactivation method, with Dunne et al demonstrated that a ≥15-minute incubation time was sufficient to inactivate a loop full of culture (Dunne, Doing et al. 2014). Other chemical inactivation reagents are used such as chloroform: methanol and chloroform: 70% ethanol which does not compromise DNA integrity (Billard-Pomares, Bleibtreu et al. 2019). Ultimately, there is no uniform consensus on the methodology of inactivating *M. tuberculosis* and therefore each method must be validated before use. We demonstrate that 70% ethanol causes a >seven-log reduction in recoverable *M. tuberculosis* in 20 minutes, removing the need for toxic solvents, heat and also allows the use of a readily available, low cost, safe, laboratory solvent-ethanol. We did not inactivate *M. abscessus* as this is a CL2 pathogen.

WGS in mycobacteria requires special consideration due to their elaborate cell wall structure, making lysis a challenge (Epperson and Strong 2020). As a result, DNA yield and quality is often unsuitable for sequencing and provides a large bottleneck to treatment and research (Kaser, Ruf et al. 2010). Mycobacterial cell walls can be disrupted via chemical or mechanical lysis.

Chemical lysis provides a lower cost method of disruption however the yield can vary between protocols and kits. Mechanical lysis involves disrupting the cell wall with beads; Aldous et al found that mechanical lysis can yield greater quantities of mycobacterial DNA than other methods (Aldous, Pounder et al. 2005). We demonstrated that bead beating leads to considerable yields in DNA downstream, but that >45 seconds beating at 10,000 rpm can degrade genomic DNA, as evidenced by two-minute bead beating with *M. tuberculosis* having DINs of <3.5 in this study. Biomass input for mycobacterial DNA extractions was also investigated in this study. We demonstrated that an O.D. of 0.1, corresponding to ∼10^7^ CFU/mL culture was the lower limit of extracting DNA that could be measured. This is still relatively high in terms of bacterial input for DNA extractions; future work will focus on optimising this process to recover DNA from lower input samples.

Targeted long read sequencing is currently advised for detecting drug resistance in TB by the WHO (World Health Organisation 2023), however in practicality Illumina platforms are often already in use in healthcare systems including low resource settings, are clinically validated and facilitate more high throughput whole genome sequencing. In addition, targeted long read sequencing cannot be used for more complex analyses including phylogenetics or transmission studies. Our method presented optimises the extraction workflow upstream of whole genome sequencing in both high and low-income settings.

## Acknowledgements and funding

Funding from the Liverpool School of Tropical Medicine Jean Clayton Fund and the Directors Catalyst Fund to DC has supported this study. This study is also funded from the UK Research Council through a PhD scholarship from the MRC Doctoral Training Partnership to CP (MR/N013514/1) and a Medical Research Council Global Challenges Fund (MR/S00467X/1) to GB. The funders had no role in study design, data collection or analysis, or the decision to submit the work for publication.

